# Noninvasive genetic surveys before and after a megafire detect displacement of migratory mule deer

**DOI:** 10.1101/2021.05.21.445205

**Authors:** Jennifer L. Brazeal, Rahel Sollmann, Benjamin N. Sacks

## Abstract

Due to climate change and past logging and fire suppression, the western US are experiencing increasingly large and frequent wildfires. Understanding how wildlife respond to these mega-fires is becoming increasingly relevant to protect and manage these populations. However, the lack of predictability inherent in such events makes studies difficult to plan. We took advantage of a large high-severity wildfire that burned adjacent to an ongoing study of mule deer (*Odocoileus hemionus*) on their summer range upslope of the fire to investigate their displacement onto our study area both immediately and upon their return to summer range the following year. We used spatial capture-recapture models in conjunction with noninvasive fecal DNA sampling to estimate density and non-spatial Pradel robust-design models to estimate apparent survival and recruitment rates. Compared to density before the fire, we observed an increase in deer density and an increase in per-capita recruitment rates one month after the fire. These findings suggest that the immediate response of at least some deer was to flee the fire upslope onto the study area rather than to downslope toward their winter range. These changes did not carry over into the following year, however, suggesting that deer formerly using the burned area as summer range may have returned there despite the high severity of the fire, or may have chosen new areas for their summer range. This suggests that, at least in the short term, the fire did not negatively affect the deer population.

## Introduction

Owing to a history of fire suppression and logging and compounded by climate change, the frequency, size, and severity of wildfires in temperate forests have increased across the Western United States (Miller et al. 2009, Westerling et al. 2011, Collins 2014, Steel et al. 2015) and globally (Williams 2013) over the past few decades. For example, just two months into the 2020 fire season, California has seen 5 of the 6 largest wildfires in the state’s 90-year recorded history (Calfire 2020). With such a dramatic rise in “mega-fires,” wildfires >10,000 ha in extent that are of higher intensity than historical wildfires (Stephens et al. 2014), it is increasingly vital to understand how forest communities respond to such large-scale disturbances. Fire can alter forest structure and the distribution of wildlife, from both direct effects arising from the fire itself, as well as from indirect effects, such as changing patterns of herbivory in response to resource availability on burned areas (Kitzberger et al. 2005, Tercero-Bucardo et al. 2007, Spitz et al. 2018).

In North American forests, mule and black-tailed deer (*Odocoileus hemionus*) are abundant and widespread. At high densities, deer have the capacity to dramatically alter forest structure and plant species through browsing pressure (Augustine and Frelich 1998, Russell et al. 2001, Rooney and Waller 2003). As such, they likely play an important role in post-megafire succession, a process that remains little understood but appears to be characterized by low conifer recruitment and high shrub cover (Collins and Roller 2013). It is therefore useful to understand how deer populations respond to wildfires in terms of abundance and distribution across the landscape. Despite the importance of this question, the lack of predictability of wildfires presents challenges to studying their impacts. A truly comprehensive understanding of how fires affect wildlife requires the ability to collect data before, during, and after the event, which is impossible to plan without advance knowledge of fire occurrence.

In the present study, we took advantage of a “natural” experiment presented by a large fire that erupted just as we were completing a noninvasive genetic density estimation study of a migratory mule deer population on their summer range in a mid-elevation mixed conifer forest in northern California (Brazeal et al. 2017). The fire burned a large portion of the population’s winter and migratory stopover ranges, directly west of their main summer range. During the summers of 2013 and 2014, we placed transects for a spatial capture-recapture (SCR) study immediately upslope of the area over which the fire would later occur. The timing of the fire presented us with the opportunity to extend the study to investigate the response of the deer population to the fire.

We investigated changes in density and population turnover on the study area before and after the King fire. Specifically, we sampled before the fire, immediately after the fire, and during the following summer to investigate hypotheses about how deer moved in response to the fire (Table 1). Based on these hypotheses, we predicted that immediately after the fire (H1-A) deer density and recruitment would remain unchanged (or decrease) as deer on the burn areas fled downslope to the winter range, effectively beginning their fall migrations early, or (H1-B) deer density and recruitment would increase as deer on the burn areas fled to adjacent areas, including onto the study area.

**Table 1.** Table of hypotheses and predictions for the response of a migratory population of mule deer (*Odocoileus hemionus*) in the El Dorado National Forest, CA, to the King Fire, which occurred September to October, 2014 adjacent to the study area. Hypotheses are at two temporal scales: immediately post-fire, from October to November, 2014 (Post-fire 1) and 1 year post-fire, in the summer of 2015 (Post-fire 2). Predictions use the following parameters: deer density (*D*), apparent survival rate (*ϕ*), and per capita recruitment rates (*f*).

| Time scale | Hypotheses | Predictions |
| --- | --- | --- |
| Post-fire 1 | **H1-A.** To escape the fire, burn-area deer moved downslope to their winter ranges. | $D_{pre-fire} \geq D_{Post-fire\ 1}$<br>$f_{pre-fire} = f_{post-fire\ 1}$<br>$\phi_{pre-fire} = \phi_{post-fire\ 1}$ |
| | **H1-B.** To escape the fire, burn-area deer moved onto our study area. | $D_{pre-fire} < D_{Post-fire\ 1}$<br>$f_{pre-fire} < f_{post-fire\ 1}$<br>$\phi_{pre-fire} = \phi_{post-fire\ 1}$ |
| Post-fire 2 | **H2-A.** Burn areas did not provide enough resources: |  |
| | **H2-A1.** Burn-area deer shifted home ranges onto our study area. | $D_{pre-fire} < D_{Post-fire\ 2}$<br>$f_{pre-fire} < f_{post-fire\ 2}$<br>$\phi_{pre-fire} \geq \phi_{post-fire\ 2}$ |
| | **H2-A2.** Burn-area deer shifted home ranges away from burn area but not onto our study area. | $D_{pre-fire} \geq D_{Post-fire\ 2}$<br>$f_{pre-fire} \geq f_{post-fire\ 2}$<br>$\phi_{pre-fire} \geq \phi_{post-fire\ 2}$ |
| | **H2-B.** Burn areas provided sufficient resources; deer returned to pre-fire home ranges | $D_{pre-fire} = D_{Post-fire\ 2}$<br>$f_{pre-fire} = f_{post-fire\ 2}$<br>$\phi_{pre-fire} = \phi_{post-fire\ 2}$ |

For the summer 1 year after the fire, we hypothesized that (H2-A) the large extent and high severity of the burn areas would result in a reduction of important resources for deer. If so, we predicted that density and recruitment the summer following the fire would either (H2-A1) increase relative to pre-fire levels, as deer originally migrating to the burn areas shifted their home ranges upslope to our study area, or (H2-A2) would remain the same or decrease if deer avoided burned areas but did not move onto our study area. We further expected apparent survival (which would only reflect emigration or deaths from the deer on our study area before the fire) to either decrease or remain the same, depending on whether or not burned areas on winter and transitional ranges impacted deer. If they did, we predicted apparent survival to decrease from direct impacts on fitness, as well as from decisions not to migrate if deer avoided moving through burned areas during spring migration.

Under the alternative hypothesis (H2-B) that burn areas did provide sufficient resources for deer, we expected deer density, recruitment, and survival to remain the same as pre-fire levels. An unavoidable limitation of this natural experiment was that if all parameters remained the same 1 year after the fire compared to before the fire, we would be unable to distinguish between hypotheses H2-B and the special case of H2-A2 where deer were not negatively impacted by burned areas on their winter and transitional ranges. We used a combination of SCR models and Pradel robust-design models to assess wildfire-associated changes in density, apparent survival and per-capita recruitment rates. Apparent survival and per-capita recruitment in these models did not differentiate between true survival and recruitment versus emigration and immigration.

## Materials and methods

### Study area

The study area was located in the Crystal Basin Recreation Area of the El Dorado National Forest in the central Sierra Nevada mountain range of Northern California (Fig. 1). The 512 km^2^ region encapsulated the critical summer range of the migratory portion of the Pacific Deer Herd mule deer population (T. Weist, California Department of Fish and Wildlife, personal communication). The elevation of the study area ranged from 1600–2300 m. Mule deer in this area migrated to their summer home ranges at elevations of 1600–2500 m. during late spring, typically between the months of April and June, and returned to their winter ranges at elevations of 600–1500 m. after the first winter storms, from mid-October to November (Hinz et al. 1981). From 13 September to 9 October, 2014, the King Fire burned 359.45 km^2^, with high vegetation burn severity (>75% canopy mortality) occurring on 198.54 km^2^ (55% of the area) (Jones et al. 2016) and high soil burn severity (<20% ground cover remaining; bare soil or ash at the surface) occurring on 89.38 km^2^ (23% of the area) (Parsons et al. 2010, USDA Forest Service, Remote Sensing Applications Center, BAER Imagery; Fig. 1). The burned areas encompassed elevations ranging 580–2100 m. The King Fire burned areas directly west and downslope of the study area, over portions of the winter and migratory stopover mule deer ranges. During our sampling years (2013–2015), the average maximum monthly temperature was 22.8°C between June and September and 13.6°C in October and November, and the average daily precipitation was 55.7 cm. (California Department of Water Resources, Station IDS: RBP, LON, VVL; 2013–2015). Water was abundant throughout the study area, including 3 large reservoirs, Ice House Reservoir (0.040 km^3^), Union Valley Reservoir (0.21 km^3^), and Loon Lake (0.052 km^3^) (California Department of Water Resources, Station IDs: ICH, UNV, LON; 2013–2015). The predominant cover types were mixed conifer forest and mixed and montane chaparral. Common plant species included a variety of conifer species, as well as shrub and browse plants, including white fir (*Abies concolor*), red fir (*Abies magnifica*), Jeffrey pine (*Pinus jeffreyi*), sugar pine (*Pinus lambertiana*), ponderosa pine (*Pinus ponderosa*), huckleberry oak (*Quercus vacciniifolia*), greenleaf manzanita (*Arctostaphylos patula*), and whitethorn ceanothus (*Ceanothus cordulatus*) (Hinz et al. 1981). Forbs and grasses were interspersed throughout the area. Land use in the study area consisted of a variety of recreational activities, including off-highway vehicle use, camping and hiking, fishing, and hunting. In addition, portions of the land were used for timber sales and grazing allotment rentals. Fire management practices at the time of the study included prescribed fire and manual thinning of vegetation.

**Fig. 1.**
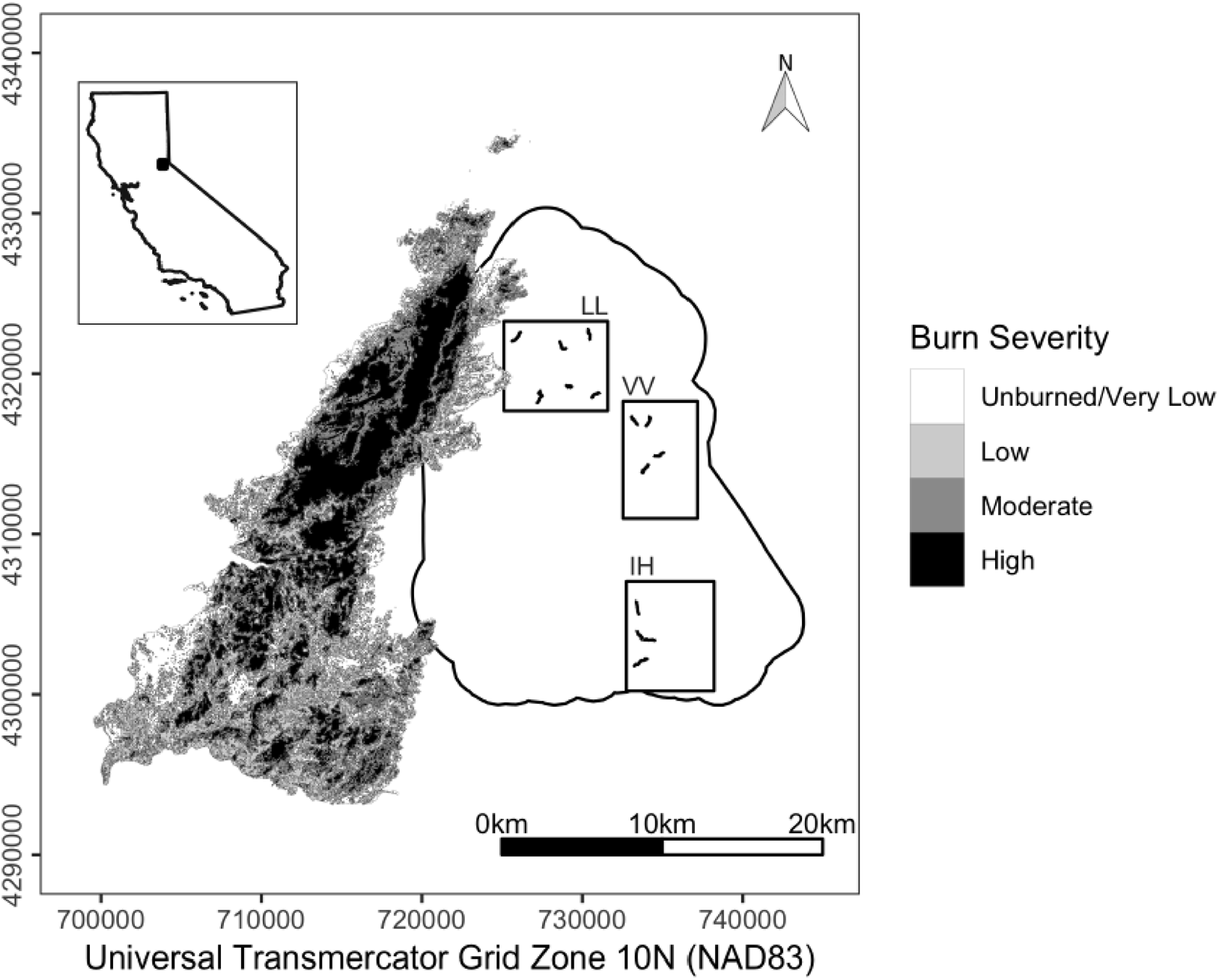
Map of the study area boundary in the El Dorado National Forest, California, with rectangles indicating the 3 sampling sites (IH = Ice House, LL = Loon Lake, VV = Van Vleck) and 1.2 km belt transects where mule deer (*Odocoileus hemionus*) pellets were collected, indicated by lines within each site (n = 14). Overlapping the study area is a surface showing the burn severity levels of the King Fire, which burned 13 September to 9 October, 2014.

### Field sampling

In 2013, we established 24 1.2 km by 2 m belt transects, randomly allocated in 4 30-km^2^ blocks throughout the study area to estimate density of a migratory population of mule deer, part of the Pacific Deer Herd, described in detail by Brazeal et al. (2017). Briefly, we selected study blocks such that no two of them shared edges and their collective habitat cover type composition was similar to the study area as a whole. Within each block, we randomly selected 6 start points for the origins of transect establishment. At each start point, we selected a random compass bearing, and established the origin at the first sign of deer (e.g., deer trail, browse, tracks). We used flagging tape to periodically mark the transect, as well as GPS points at 100 m intervals. We followed deer trails when present, to optimize capture and recapture rates (Brinkman et al. 2011, Lounsberry et al. 2015, Brazeal et al. 2017; Furnas et al. 2020), and followed the original random compass bearing when the trail was lost, until we established a transect 1–1.2 km in length.

For the present study, we used a subset of 14 transects from the original 24, from 3 of the 4 original study blocks (Fig. 1). We could not sample from one of the original study blocks during Post-fire1 because of road closures associated with the fire. For all analyses, we included samples only from the 14 transects we were able to sample in all sampling periods. We collected samples during 2013–2015 during four discrete sampling periods. Two of the sampling periods occurred before the King fire (3 June, 2013 to 24 July, 2013 and 23 June, 2014 to 10 September, 2014), one occurred immediately post-fire (Post-fire1; 5 October, 2014 to 10 November, 2014), and one occurred 1 year post-fire (Post-fire2; 13 July, 2015 to 31 August, 2015). During each sampling period, we visited and collected pellets on at least 3 sampling occasions per transect. In the 2014 pre-fire sampling period, we visited transects 3–6 times. We visited each transect every 7–10 days to ensure enough time for pellets to accumulate while limiting the amount of time they were exposed to the environment. On each visit, we scanned for and collected pellets 2 m on either side from the transect line. To minimize collection of samples that were unlikely to yield DNA, we only sampled pellets that showed no obvious signs of degradation on the pellet surface (i.e., no cracks and not dull in color). We collected 4–6 pellets from each distinct pile and stored them in 95–100% ethanol in 15 mL centrifuge tubes.

We completed laboratory analyses at the Mammalian Ecology and Conservation Unit of the University of California Davis Veterinary Genetics Laboratory, following protocols described by Brazeal et al. (2017). Briefly, we genotyped loci in a single multiplex assay using 10 microsatellite loci (ADCYC, BM5606, Celb9, Cervid1, ETH152, SBTD04, SBTD05, SBTD06, SBTD07, TGLA94) and SRY, a sex marker from a highly conserved region on the Y chromosome (Lounsberry et al. 2015). We manually binned all genotype allele sizes and used replicate samples to construct consensus multi-locus genotypes. Because of inconsistent amplification rates during post-fire periods, we eliminated locus SBTD04 from all downstream analyses.

For individual identification, we used only samples that amplified successfully at 7 of the remaining 9 loci to cluster genotypes into individual IDs based on an optimal maximum threshold number of mismatching alleles (Galpern et al. 2012). This procedure was described in detail previously, as was estimation of the allelic dropout rate (<1.2%; Brazeal et al. 2017). The assignment criteria in Brazeal et al. (2017), combined with the genotyping error rate, resulted in an estimated assignment accuracy rate of ≥97%. Once we determined the optimal number of mismatching alleles for the set of reduced autosomal loci in this study (9 loci versus 10 loci), we manually determined the assignment of any multi-locus genotypes that were matched to more than one individual identity using a set of identification rules. First, we eliminated a potential match if any of the mismatches of the focal genotype to the potential match could not be due to simple allelic dropout. Second, we compared the sex of the focal genotype to its potential matches, and eliminated potential matches whose sex did not match. Finally, among remaining potential matches, those from the same transect were presumed to be the same individual and those from different transects were presumed to be different individuals. As a standard metric of discriminatory value of the assay, we also estimated the probability that two siblings shared the same multilocus genotype (PID_*SIB*_; Wilberg and Dreher 2004).

### Data summarization

Before conducting capture-recapture analyses, we investigated numbers of individuals detected per transect as an index of abundance across sampling periods and transect visits (which we hereafter refer to as sampling “occasions”) within sampling periods. This method did not account for imperfect or varying detection probabilities, but served as a relative index to gauge temporal effects on deer detections. We performed Poisson regressions of the average count of individuals per transect (response variable) on sampling period and occasion (predictor variables). We treated sampling period as a categorical variable with three levels: pre-fire (i.e., 2013 and 2014, pooled), Post-fire1, and Post-fire2, and occasion as a continuous variable to represent a temporal trend. We examined both main effects and interaction terms. We selected the model with lowest AIC_*c*_ as the most parsimonious model (Burnham and Anderson 2002) and used Wald tests to determine the significance of beta parameters.

### Density

To estimate density, we used multi-session closed Huggins SCR models (Huggins 1989, Borchers and Efford 2008). In multi-session SCR models, a “session” is a survey that is considered independent of other sessions, either spatially or temporally, but parameters may be estimated using data across sessions (Efford 2018). Our sampling periods corresponded to these sessions. The population was assumed to be closed demographically within each session and to test the assumption of closure within sessions, we used a closure test that was robust to individual heterogeneity in detection probability (Otis et al. 1978), as well as tests that were robust to variation in detection probability over sampling occasions (Stanley and Burnham 1999). The SCR models consist of state and observation model components. The state model represents the distribution of animal home range centers across space and the observation model describes the probability of detecting an animal as a declining function of distance between the animal’s home range center and a given detector. For detectors, we discretized each of the 14 transects to multiple point detectors, spaced 200 m apart, though because transects were not perfectly linear, Euclidean distances between detector points were usually <200 m. This procedure resulted in a total of 75 detectors and an average of 5.36 (SD = 0.84) detectors per transect. We assigned each detection of an individual to its closest detector and treated multiple detections of an individual at a detector during the same occasion as a single detection (corresponding to the “proximity detector” type in secr) to avoid double-counting a single defecation event.

For the state model component, we assumed a homogeneous Poisson point process distribution of home range centers. For the observation model component, we estimated the expected probability of detection of an animal whose home range was centered at a given detector (*λ*_*O*_) and the scale of movement of the animal (*σ*; a scaling factor representing the decline in detection as distance increases from a detector, which biologically represents a model-specific, relative measure of an average home range radius), using a hazard exponential detection function:

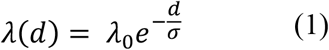

where *λ(d)* refers to the expected probability of detection for an individual whose home range center lies a distance (*d*) from a given detector. We parameterized the detection function over a discrete region of integration, which we refer to as the habitat mask. To make the habitat mask, we buffered each transect by 2 km on all sides, which we estimated to be large enough to contain all animals with home ranges overlapping our point detectors, and removed areas covered by water. We used a 2-class mixture model with known class memberships to model sex differences in detection probability and to obtain sex-specific estimates of *λ*_*O*_ and *σ* (Borchers and Efford 2008, Pledger and Phillpot 2008). This method also provided estimates of the mixture proportions of the 2 classes (*pmix*), which we used to estimate the sex ratio of the population. Specifically, we estimated the female to male (F:M) sex ratio as the ratio of the estimates of female and male mixture proportions, and used the delta method to estimate the corresponding standard error (Powell 2007). We used the conditional likelihood, which conditioned on the detected individuals, and estimated *D* as a derived parameter using a Horvitz-Thompson-like estimator (Borchers and Efford 2008). We tested models that held *λ*_*O*_ constant or included combinations of the following covariates: sex (i.e., the 2-class mixture factor), session, and occasion (assuming a linear relationship). We held *σ* constant across sessions but allowed it to vary by sex. We considered models with a mixture proportion of sexes either constant or variable across sessions. Because the number of occasions varied across transects in 2014, we included a binary usage matrix indicating whether or not a detector was sampled on any given occasion. We implemented these models in the R package secr (Efford 2018) run in R package version 3.1.5., https://CRAN.R-project.org/package=secr, accessed 15 April, 2018.

### Per-capita Recruitment and Apparent Survival

We estimated per-capita recruitment (*f*) and apparent survival (*ϕ)* using robust-design Pradel recruitment models (Pollock 1982, Pradel 1996). Robust-design models involve two temporal levels of sampling. Multiple secondary periods are nested within primary periods (corresponding to occasions and sessions in our SCR application, respectively). Within a primary period, the population is assumed to be demographically closed. Between primary periods, the population is open to births, deaths, immigration, and emigration. The secondary periods enabled robust estimation of detection probability from repeated sampling of the population over a closed period, while the primary periods allowed for estimation of apparent survival rates (from both deaths and emigration) and per-capita recruitment rates (from births and immigration). We will hereafter refer to sessions or primary periods as “sampling periods” and occasions or secondary periods as “sampling occasions” when we are not referring to a specific model. Otherwise, we will maintain the appropriate naming for a given model type.

We only considered captures from 2014 pre-fire on secondary periods 1–3, as some transects were not sampled beyond secondary period 3. Because individuals were sometimes detected on multiple transects, we collapsed encounter histories across all transects. We elected to reduce the number of secondary periods in 2014 pre-fire to the same number as in the post-fire primary periods (i.e., 3 secondary periods each). We considered the time interval between 2014 pre-fire and Post-fire1 to be 1, which would approximate a 1-month time interval. Between 2013 and 2014, as well as Post-fire1 and Post-fire2, we specified 10 time intervals, reflecting the approximately 10 months separating the end of one primary period and start of the next. We estimated population turnover rates, represented as per-capita recruitment 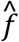 and survival 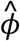, on the specified time interval scale. For 2014 pre-fire, we performed closure tests as described above, with a subset of only the first 3 sampling occasions.

We tested a total of 70 models, considering sex and primary period as possible predictors of 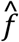and 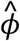. Because adult male deer were hunted annually on the study area, we only considered models in which 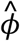 varied by sex or sex and primary period. On the other hand, we had no *a priori* expectations regarding variation in 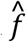. Therefore, we considered models where 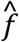 was constant, varied with sex, or varied with sex and primary period (both with and without an interaction effect).

We held detection probability (*p*) constant or allowed it to vary by sex, primary period, secondary period (assuming a linear relationship), and combinations of these variables. Our sampling was asynchronous across transects, so the traditional *M*_*t*_ model of Otis et al. (1978), where *p* varies by secondary period, was inappropriate. However, by testing for a linear effect of secondary period on detection probability, we intended to account for a potentially higher detection rate during the initial versus later periods because we did not clear transects before our first collection of pellets. That is, the first visits may have had higher detection probabilities because we collected pellets older than 7 to 10 days old, leading to a possible decrease in *p* as the number of visits to a transect increased. We only considered models where the number of deer undetected in each period (*f*0) varied by both sex and primary period. We used AIC_*c*_ to select the most parsimonious models and only considered those that were ≤2 *Δ*AIC_*c*_ from the top-ranking model (Burnham and Anderson 2002, Arnold 2010).

## Results

We collected 881 fecal pellet samples during 2013 (n = 241), 2014 pre-fire (n = 265), Post-fire1 (n = 213), and Post-fire2 (n = 162; Table 2). Of those samples, 497 (56%) amplified at 7 of the 9 loci used. We estimated the probability of two siblings sharing the same multilocus genotype (*PID*_*SIB*_) at 5.9 *⨯* 10^*-4*^. Our optimal threshold for allelic mismatches was 5 (28%, Fig. 2). In total, we identified 202 individuals from the 497 genotypes. After condensing multiple pellet samples of a given individual at the same detector on the same transect visit into a single detection, we retained 445 detections of the 202 individuals. For the robust-design models (see below), we removed secondary occasions 4–6 from the pre-fire 2014 primary period, retaining 57 of the 80 individuals detected during the pre-fire 2014 period.

**Table 2.** Total number of fecal samples collected from 14 transects in the El Dorado National Forest, CA for non-invasive genetic abundance-estimation in a population of migratory mule deer (*Odocoileus hemionus*). Shown for each sampling period are the number (No.) of deer pellet group samples collected, the number of these samples that amplified at ≥7 loci and were used for individual identification (No. genotypes), the number of these genotypes that were independent, i.e., after removing genotypes corresponding to individuals recaptured at the same detector and on the same occasion, the number of individuals detected each period, and the number of these individuals captured for the first time in that period.

| Sampling Period | No.<br>samples | No.<br>genotypes | No.<br>independent<br>genotypes | No. Individuals<sup>†</sup><br>(F/M) | No. first captures<br>(F/M) |
| --- | --- | --- | --- | --- | --- |
| Pre-fire 2013 | 241 | 99 | 89 | 65 (43/22) | 65 (43/22) |
| Pre-fire 2014 | 265 | 165 | 148 | 80 (53/27) | 62 (37/25) |
| Post-fire 2014 | 213 | 128 | 115 | 77 (54/23) | 44 (25/19) |
| Post-fire 2015 | 162 | 105 | 93 | 66 (46/20) | 31 (17/14) |
| Totals | 881 | 497 | 445 | -- | 202 (122/80) |

<sup>†</sup>The total number of individuals includes both individuals newly captured and individuals captured during previous sampling periods.

**Fig. 2.**
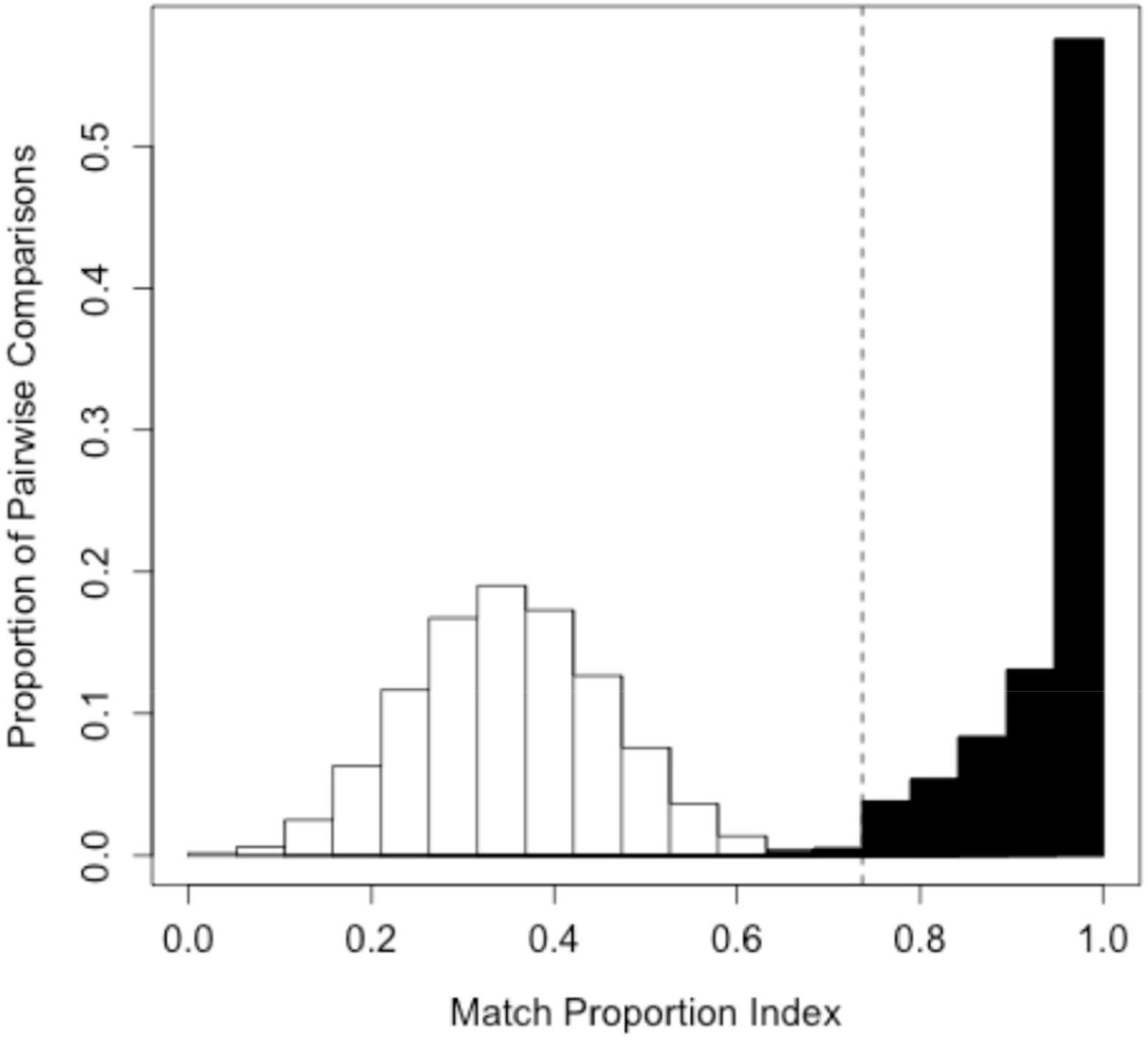
Plot of two histograms showing the match proportion index estimated from R package allelematch (Galpern et al. 2012). The first histogram (white bars) show the level of similarity in pairwise comparisons between distinct individuals. The second histogram (black bars) shows the match proportion indices between genotypes attributed to the same individual. We identified individuals from multilocus microsatellite genotype sequences (9 loci and a sex marker) using an optimal mismatch threshold of 5 alleles. This threshold corresponds to the dashed vertical line on the plot. The individuals were mule deer (*Odocoileus hemionus*) detected from noninvasive fecal genetic surveys in the El Dorado National Forest in Northern CA, USA between 2013 and 2015.

### Numbers of individuals detected

In Post-fire1, there was an increase in the average number of individuals detected on transects compared to pre-fire sampling periods (Fig. 3). However, this trend was only evident in the first and second sampling occasions, with a steep decline on the third sampling occasion.

**Fig. 3.**
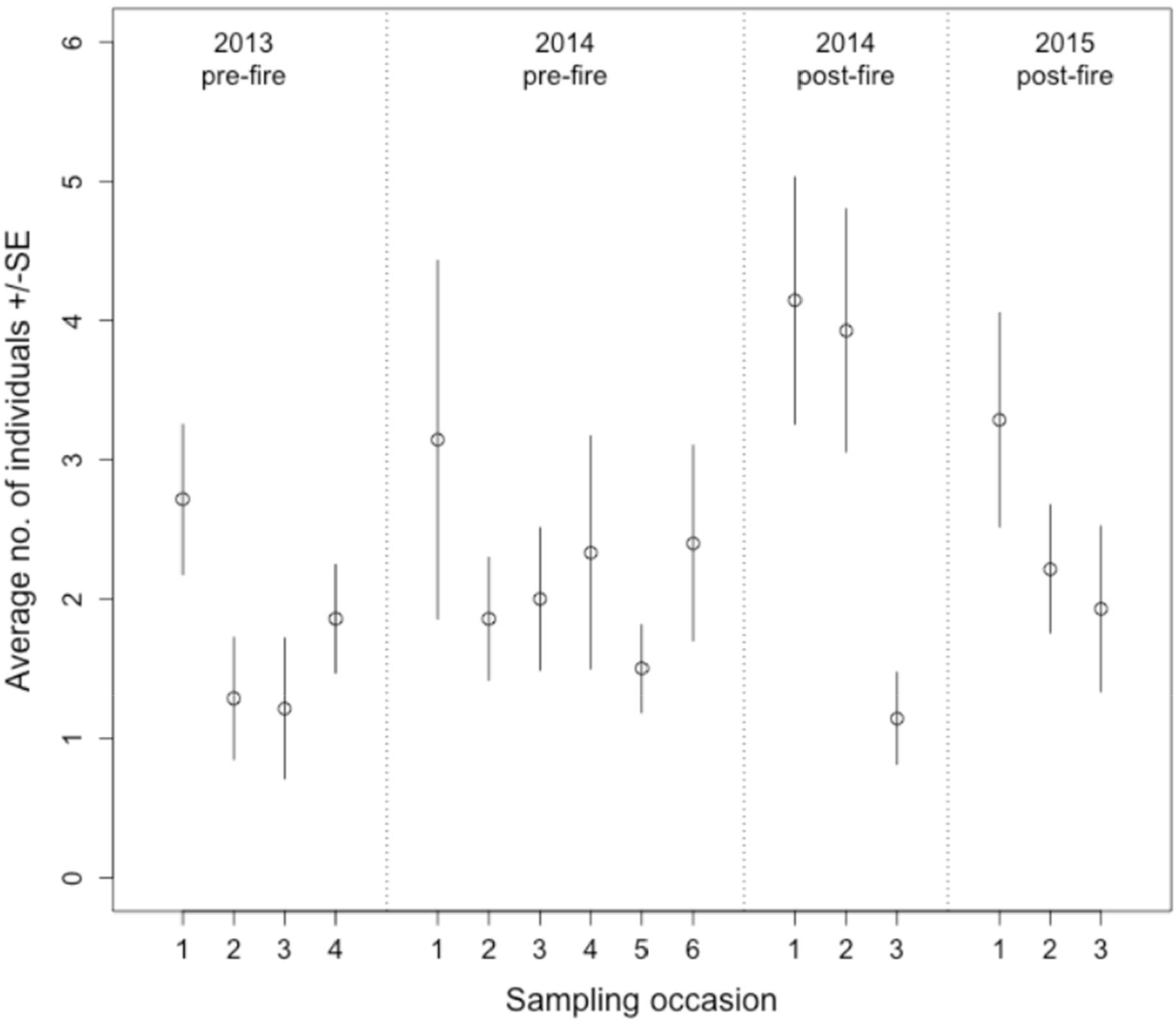
Average number of individual mule deer (*Odocoileus hemionus*) detected per transect (n = 14 transects), based on distinct multilocus genotypes, with standard error bars in the El Dorado National Forest, California, across 4 sampling periods: summer 2013 (2013 pre-fire), summer 2014 (2014 pre-fire), immediately post-fire in October and November, 2014 (2014 post-fire), and summer 2015 (2015 post-fire). Within each period, we show the average number of individuals detected on each sampling occasion, which represent the sequential visits to each transect. On all occasions, we sampled 14 transects, with the exception of 2014 pre-fire, in which we sampled 14 transects on occasions 1–3, 12 transects on occasions 4–5, and 10 transects on occasion 6.

Correspondingly, the Poisson regression model with the lowest AIC_c_ indicated that the average number of individuals detected was higher during Post-fire1 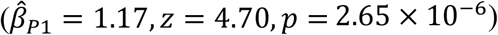 and, to a lesser extent, Post-fire2 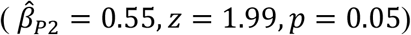 than the two pre-fire sampling periods. The interaction between period and occasion was also significant, indicating that although numbers of individuals detected declined over the course of all sampling periods, the magnitude of this decline was significantly greater during Post-fire1 than the other sampling periods 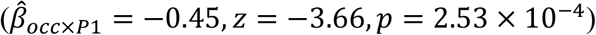.

### Tests of Closure within Sampling Periods

Based on the Otis et al. (1978) test, we detected no significant violations of the closure assumption in either pre-fire sampling period or in Post-fire2, but detected a significant violation of the closure assumption in Post-fire1 (*z = -*2.45, *p =* 7.15 *⨯* 10^*-3*^). The two Stanley and Burnham (1999) tests relevant to recruitment to Post-fire1 also were significant (NR vs. Jolly-Seber 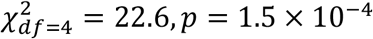 and *M*_*t*_ vs. NM 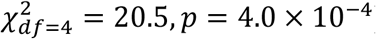). The relatively strong decline in detections in the third sampling occasion of Post-fire1 (Fig. 3) suggested that violations in closure within Post-fire1 likely reflected the start of winter migrations off the study area. To remove any bias associated with these migrations, we pooled the second and third occasions in Post-fire1 to obtain unbiased estimates of the number of individuals present in the initial sampling period, as recommended by Kendall (1999).

### Spatial capture-recapture

The SCR model with the lowest AIC_c_ included sex and occasion as predictors of *λ*_*O*_, and sex as a predictor of *σ*. The only other model ≤2 *Δ*AIC_c_ was fully nested within the top model, with no effect of sex on 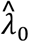 and 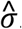. We present results for the top model only (Table 3). The expected encounter rate declined over successive occasions 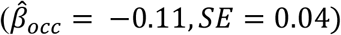. Females had a higher expected encounter rate 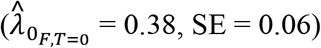 than males 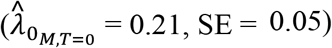, while males had a larger scale of movement (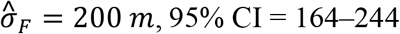 and 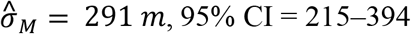). The estimate of *D* increased immediately post-fire (i.e., from 2014 pre-fire to Post-fire1), though its 95% CI in Post-fire1 overlapped with pre-fire estimates. In Post-fire2, 1 year after the fire, 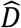 was also non-significantly higher than pre-fire levels, though less so. Estimates of *D* ranged from 4.1−4.4 deer/km^2^ in pre-fire periods and 4.9−5.6 deer/km^2^ in post-fire periods. The F:M sex ratio estimated from this model was 2.82 (95% CI = 2.54–3.10).

**Table 3.**
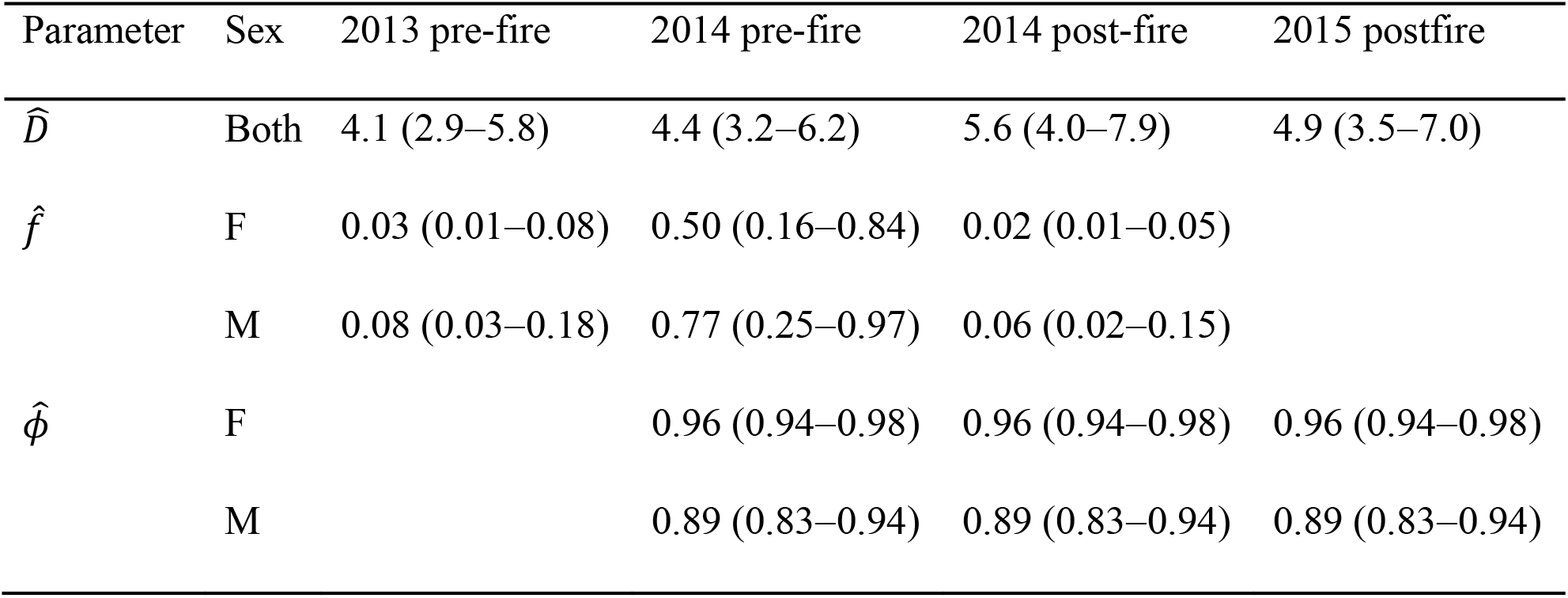
Estimates of mule deer (*Odocoileus hemionus*) density (*D*), per-capita recruitment rates (*f*), and apparent survival rates (*ϕ*), before (2013 pre-fire and 2014 pre-fire) and after (2014 post-fire and 2015 post-fire) the King Fire burned areas adjacent to the deer’s summer range in the El Dorado National Forest, CA. Numbers in parentheses are 95% confidence intervals. Estimate of *D* are pooled for both sexes (Both) and obtained with spatial capture-recapture models, while estimates of *f* and *ϕ* are specific to females (F) and males (M) and obtained with non-spatial robust design capture-recapture models. Estimates of *f*_*i*_ correspond to the per-capita rate of additions to the population between sampling periods *i* and *i* + 1. Estimates of *ϕ*_*i*_ correspond to survival rates between periods *i* – 1 and *i*.

### Robust-design recruitment and survival estimates

The Pradel robust-design model with the lowest AIC_c_ contained sex as a predictor of *ϕ*, sex and primary period as predictors of *f*, and sex and secondary period as predictors of *p*. The only other model ≤2 *Δ*AIC_c_ (*Δ*AIC_c_ = 1.13) was fully nested within the top model, with the effect of sex on *f* removed. We only present results from the top model. Detection probability declined over secondary periods within primary periods 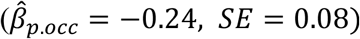. The monthly per-capita recruitment rates for both sexes was significantly higher for 2014 pre-fire than for other primary periods, suggesting a larger number of additions to the population between 2014 pre-fire and Post-fire1 than between other consecutive primary periods (Table 3). The Post-fire1 monthly per capita recruitment rate was similar to the estimate for 2013, indicating a similar rate of addition to the population between the two post-fire periods and the two pre-fire periods. Monthly survival 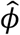 was lower for males than females (Table 3). The trend in abundance 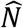 from the robust-design models was similar to the trend in 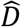 from the SCR models, except the confidence intervals for female 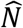 in Post-fire1 did not overlap primary period estimates, indicating a significant increase in estimated abundance immediately after the fire in Post-fire1 (Fig. 4).

**Fig. 4.**
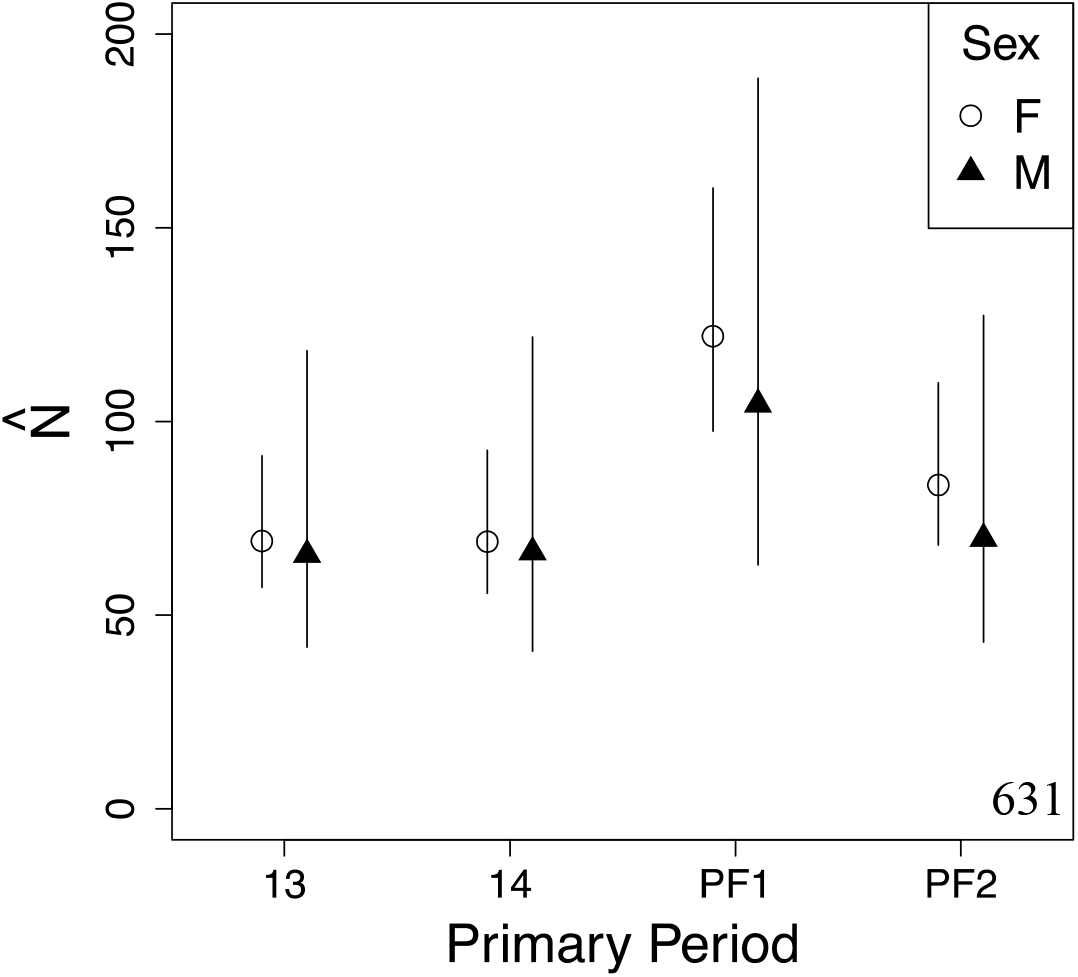
Plot of sex-specific abundance estimates, 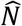, for a population of mule deer (*Odocoileus hemionus*) in the El Dorado National Forest, CA before and after a large wildfire burned areas west and downslope of the study area. Estimates were obtained with a robust design Pradel capture-recapture model fit to noninvasive genetic detection data and are shown with 95% confidence intervals. Results are reported for each primary period, during which 14 transects were visited 3 times each. We sampled in the summer of 2013 (13), the summer of 2014 (14), immediately post-fire, in October and November, 2014 (PF1), and 1 year post-fire in the summer of 2015 (PF2).

## Discussion

We assessed the response of a migratory population of mule deer to a mega-fire in the Central Sierra Nevada, California. We found that there were significant additions to the population of deer on our fire-adjacent study area immediately post-fire, which could only be attributed to immigration, as births could not have occurred during the months of September and October. Per capita-recruitment rates significantly increased immediately post-fire, with a corresponding (non-significant) increase in density. These findings supported the hypothesis that deer were displaced by the fire onto our study area immediately post-fire (hypothesis H1-A1, Table 1). Although a general increase in movement of deer beginning their winter migrations likely affected our results to some extent, it is unlikely to have explained the observed trends. In particular, closure tests suggested there was recruitment in the 1-month interval between 2014 pre-fire and Post-fire1, but not within Post-fire1, indicating the observed additions to the population immediately after the fire mostly occurred before sampling for that period began.

One year after the fire, no population parameter estimates differed significantly from the pre-fire estimates, suggesting that the increase in the per capita recruitment rate and density after the fire was due to immediate and temporary displacement of individuals from the burned area.

Conversely, we did not see a decrease in recruitment, survival, or density one year post fire, suggesting that deer resident on the study area did not suffer significant losses during their migrations through the burn area or on their winter range from lack of cover or forage. These results align with predictions for hypotheses H2-A2 and H2-B (Table 1). It is possible that the burn areas provided sufficient cover and forage for the deer whose home ranges were within the burn perimeter pre-fire (H2-B). Prior studies have suggested that mule deer generally are resilient to and may even benefit from wildfire. The increase in heterogeneity of forest structure resulting from wildfire can provide an increase in forage quality and quantity in open areas, while retaining adjacent vegetative cover (Hobbs and Spowart 1984, Carlson et al. 1993, Long et al. 2008). These effects, however, are dependent on fire size and severity (Wan 2014), as well as time since fire (Carlson et al. 1993). Given that a pattern of large, contiguous high-severity burn areas was particularly characteristic of the King Fire (Stevens et al. 2017) and one year provides limited time for forage regrowth in these severely affected areas, H2-B seems less likely (though deer were found to use all burn severities equally three years post-fire; RS, unpublished data). Alternatively, deer initially displaced by the fire might simply have summered on their winter range or somewhere other than our study area the following year (H2-A2), including shifting to areas within the fire footprint that suffered lower levels of disturbance. Studies in which individual movements are tracked through telemetry before, during, and after a high severity fire would provide more direct data to address such questions.

It is also important to note that our study was limited to the migratory deer with summer home ranges away from the burn area, and that we considered responses at relatively short time scales. We did not have information on deer within the burn perimeter before or after the fire, so their movement patterns, survival rates, and recruitment rates were unknown. In addition, there is potential for a time lag in the response of mule deer populations to environmental conditions, as maternal condition can affect subsequent recruitment and the survival or fitness of young (e.g., Monteith et al. 2009, Ciuti et al. 2015). Future studies exploring the effects of mega-fires on deer with home ranges on the burn area, and longer-term studies that might reveal effects on survival and recruitment occurring at longer time scales than 1 year would help elucidate the response of mule deer to mega-fires.

The combination of non-invasive genetic sampling and SCR methods are increasingly used to estimate abundance and density of mule deer populations in Western North America (e.g., Brinkman et al. 2011, Lounsberry et al. 2014, Brazeal et al. 2017) and most recently have been employed at multiregional scales conducive to coordinated state or provincial-scale wildlife management (Furnas et al. 2018, 2020). In addition, open and robust-design SCR models have recently become available (Gardner et al. 2010, Ergon and Gardner 2014, Royle et al. 2016, Gardner et al. 2018) and future studies might benefit from employing a fully integrated spatial approach to estimate density and other demographic parameters. With projected increases in large-scale disturbance events related to climate change (Spracklen et al. 2009, Westerling et al. 2011, Barbero et al. 2015), including mega-fires such as the King Fire, these methods offer promising ways to increase our understanding of species responses to such events.

## Acknowledgments

We thank T. Weist, S. D. Blair, P. C. Keating and R. W. Strabel for logistical support and field assistance field work, Z. T. Lounsberry for laboratory oversight and training, and F. A. Calderon and T. J. Kalani for laboratory analyses. Sierra Pacific Industries provided access to their lands on the study area. R. A. Baldwin, G. Kelley, and N. H. provided helpful comments and statistical. Extension of a project initially funded by the California Department of Fish and Wildlife, Big Game Management Account (Grant No. NC-8001-11) was funded by the conservation fund of the Mammalian Ecology and Conservation Unit of the Veterinary Genetics Laboratory at the University of California Davis.

